# Climate and regional plant richness drive diet specialization in butterfly caterpillars

**DOI:** 10.1101/2025.01.10.632438

**Authors:** Collin P. Gross, Akito Y. Kawahara, Barnabas H. Daru

## Abstract

Studies of coevolution, ecosystem processes, and latitudinal diversity gradients are improved by understanding variation in resource specialization. Insect herbivory is one of the most ubiquitous terrestrial ecological associations that drives the evolution of plants and insects. However, a broad understanding of how and why herbivore diet specificity varies worldwide is lacking. Here, we use global datasets of butterfly and plant distributions to investigate patterns and drivers of butterfly larval diet breadth. Hostplant richness and phylogenetic diversity showed non-monotonic latitudinal patterns that narrowed near the equator and temperate latitudes and broadened at mid-tropical latitudes and the poles. Diet breadth showed a negative relationship with plant family richness, but we also uncovered an interaction between precipitation seasonality and temperature. Our study builds on 60 years of study of butterfly-hostplant evolution to provide valuable insights into how these processes continue to shape plant and herbivore dynamics in response to global environmental changes.

## Introduction

Variation in the magnitude, direction, and breadth of interspecific interactions influences multiple ecological and evolutionary processes. Differences in dietary specialization have received attention for their effects on organisms, ecological communities, and ecosystems at multiple spatial, temporal, and taxonomic scales. Variation in specialization can affect resource competition and coexistence (Barnes & Murphy 2023; Becker *et al*. 2022), species extinction risks and response to disturbance (Devictor *et al*. 2008; Petrocelli *et al*. 2024; Slatyer *et al*.

2013), the stability of interaction networks (Mougi & Kondoh 2012; Zografou *et al*. 2020), and top-down effects on primary productivity (Singer *et al*. 2014). At broad scales, more intense and specialized interactions near the equator relative to the poles may influence the latitudinal diversity gradient – in low-latitude tropical habitats, ecological specialization can allow for “niche packing” of more species with narrow niches – but specialization itself may also evolve in response to intense competition from greater number of species at lower latitudes (Dyer & Forister 2019; Hargreaves 2024; Schemske *et al*. 2009). Generalism may facilitate species range expansion (Slatyer *et al*. 2013), and variation in diversification rates among lineages can be attributed to differences in dietary specialization (Day *et al*. 2016; Hardy & Otto 2014).

However, fundamental questions about how diet breadth varies at a global scale, such as whether diet breadth mirrors latitudinal gradients in resource richness (Forister *et al*. 2015), or which climatic factors influence diet breadth among species and assemblages, remain relatively unexplored.

Insect herbivory is one of the most ubiquitous and consequential species interactions in terrestrial ecosystems worldwide (Seastedt & Crossley 1984; Yang & Gratton 2014). Unlike terrestrial vertebrate herbivores, herbivorous insects are typically found in direct association with the plants they feed on at one or more stages of their life cycle, and in many cases are best classified as parasites (Agosta *et al*. 2010; Agrawal & Hastings 2023; Yotoko *et al*. 2005). Because of this close association, herbivorous insects are collectively known for their narrow diet breadth, the result of reciprocal selection and coadaptation that pushes herbivorous insects to specialize on plants with unique chemical defenses and discourages host-switching (Braga *et al*. 2021; Janz & Nylin 1998; van der Linden *et al*. 2021; Nishida 2002).

Theoretical work predicts a negative relationship between resource diversity and consumer diet breadth when generalism carries a fitness cost (Ackermann & Doebeli 2004; Bruijning *et al*. 2024), generating adaptive radiations of consumers into multiple specialized niches. Thus, lower latitudes are hypothesized to harbor more specialized insect herbivores because of the diversity of plant hosts, while the most speciose plant lineages allow for the evolution of a greater number of specialists (Dyer & Forister 2019; Forister *et al*. 2015).

Insect lineages colonizing islands with unexploited plant resources may be driven towards specialization over evolutionary time (Gillespie & Roderick 2002; Hembry *et al*. 2021; Losos & Ricklefs 2009). On the other hand, islands are limited in their plant species pools by size and distance from mainland colonization sources (Borregaard *et al*. 2016; MacArthur & Wilson 1967), perhaps limiting the evolution of specialized herbivore diets. Which of these patterns prevails in island habitats is likely a function of the insect assemblages on islands: assemblages composed of largely dispersal-limited island-endemic species are likely to be more specialized on island plants, while assemblages predominantly composed of wide-ranging species whose range includes islands will likely be more generalist on average (Hurtado *et al*. 2024; Lancaster 2020).

A possible proximal evolutionary explanation for global geographic variation in insect diet breadth is variation in plant species richness, which represents a range of discretely usable resources that can promote diversification and specialization (Braga *et al*. 2021; Ehrlich & Raven 1964; Fordyce 2010). Yet, plant species richness varies along global climate gradients, which themselves may directly impact insects (Cai *et al*. 2023; Daru 2024b). For example, highly seasonal climates may limit plant growth and therefore herbivore resource availability, but these climates also limit insect flight and feeding periods due to physiological tolerances. For herbivorous insects in seasonal (e.g. high-latitude) environments, climate itself may limit their ability to feed and reach sexual maturity on any hostplant, or the time to search for a suitable living hostplant is limited by host climate tolerance (Augustine & Kingsolver 2018; Nylin 1988). Where insect and hostplant climate niches differ, we might expect selection for broader insect diets in seasonal habitats as search time for the “correct” hostplant becomes costly (Neu *et al*. 2021). Alternatively, if several hostplants are available, insects in seasonal habitats may instead opt to specialize on hosts that maximize growth and minimize development time (Nylin 1988; Papaj *et al*. 2007).

Here we focus our investigation on plant-feeding butterfly caterpillars (Papilionoidea), for which the ecological and evolutionary consequences of dietary specialization are particularly well-studied (Ehrlich & Raven 1964; Fordyce 2010; Forister *et al*. 2015; Janz & Nylin 1998).

Butterflies and plants have been a fundamental system for the study of coevolutionary relationships since Ehrlich and Raven’s foundational 1964 study of the evolutionary arms race between chemical-resistant caterpillars and their chemically-defended hosts. These authors posited that reciprocal selection acting on herbivores and plants helped explain both the striking diversity of specialist butterflies as well as the secondary metabolites of many plant taxa. In the 60 years since this publication, numerous studies have tested Ehrlich and Raven’s hypotheses of coevolution and diversification (e.g. Hardy & Otto 2014; Janz & Nylin 1998; van der Linden *et al*. 2021), but open questions remain about the geographic context and patterns that might follow from these mechanisms.

We were specifically interested in how climatic, habitat, and resource-availability variables affected mean diet breadth of local butterfly assemblages and individual species, as well as how island-endemic species differ from mainland species in their diet breadth. While similar analyses of global patterns in diet breadth have been published (e.g., Forister *et al*. 2015), they have largely relied on geographically-biased point observations, have not explicitly incorporated phylogenetic information for both insects and their hosts, and did not explicitly address island assemblages. Our approach uses detailed range maps for both butterfly and plant species (Daru 2024a, b), comprehensive phylogenies (Jin & Qian 2022; Kawahara *et al*. 2023), and information on hostplant associations (Kawahara *et al*. 2023; Robinson *et al*. 2023; Shirey *et al*. 2022) to examine global patterns and putative drivers of diet breadth. We hypothesize that diet breadth will show a negative relationship with latitude, with more specialized species in regions with greater plant diversity; and that species on islands are more specialized than their mainland relatives.

## Methods

*Caterpillar and plant distributions.* We used comprehensive species-level native distribution maps for 10,372 species of butterflies of the world and 151,508 species of plants (Daru 2024b, a). Briefly, these distribution maps were modeled based on point data available from the Global Biodiversity Information Facility (GBIF 2023), clade-specific constraints on dispersal (Louca 2021), and species distribution models using maximum entropy. Each species range map was stacked at the resolution of 100 km × 100 km grid cells to generate a presence-absence matrix for species in 13,316 grid cells using the *polys2comm* function in the R package phyloregion (Daru *et al*. 2020).

*Caterpillar diets.* We characterized caterpillar hostplants at the family level, which has been adopted as the standard taxonomic rank for examining host use evolution (Braga *et al*. 2020; Kawahara *et al*. 2023). These data were obtained from multiple sources, including LepTraits (Shirey *et al*. 2022), the NHM HOSTS database (Robinson *et al*. 2023), and Kawahara *et al*. (2023), and filtered to retain only those for which at least two records were documented.

Because closely related hostplant families may share similar profiles of secondary metabolites, caterpillars may be more likely to feed on more closely related families (Ehrlich & Raven 1964; van der Linden *et al*. 2021). Thus, in addition to examining the number of hostplant families used by caterpillar species, we also used a family-level phylogeny of plants based on the supertree by Jin & Qian (2022) to inform diet breadth. Specifically, we measured hostplant phylogenetic diversity (PD) as the total branch lengths shared by all host families (Faith 1992), as well as the standard effect size (SES) of mean pairwise distance (MPD) and mean nearest taxonomic distance (MNTD) between hostplant families (Sessa *et al*. 2018). These latter metrics represent measures of how similar a caterpillar’s assemblage of hostplants is relative to a null distribution of random families drawn from the phylogeny, where negative values represent families that are more closely related than a null expectation (clustered), and positive values represent families that are more distantly related than random (overdispersed). We calculated SES_MPD_ and SES_MNTD_ using the R package picante (Kembel *et al*. 2010), using the *taxa.labels* algorithm to generate null distributions. When caterpillars were only documented to feed on one hostplant family, we included the most distant tips of the tree for the family as a way of measuring the phylogenetic distances within the family. For simplicity we group our four measures of diet breadth into two categories: “diet richness” for the count of hostplants and hostplant phylogenetic diversity, and “diet dispersion” for SES_MPD_ and SES_MNTD_. The general term “diet breadth” as used throughout this manuscript includes all four metrics.

*Predictor variables.* To examine biogeographic drivers of caterpillar diet breadth, we used two separate sets of predictors. First, we used the 19 bioclimatic variables from WorldClim (Fick & Hijmans 2017); elevation, net primary productivity, percent tree cover, non-tree vegetation cover, and unvegetated cover from MODIS (Dubayah *et al*. 2021; Running *et al*. 2021; Townsend & DiMiceli 2015); and the richness of all plant families, species richness of hostplant families for local caterpillars, and total species richness of plants as our initial pool to investigate the role of climate, habitat, and host availability on diet breadth. For climate, elevation, and vegetation we used the value corresponding to the centroid of each grid cell at a resolution of 10 minutes of a degree, while for plant richness, we summed across all species or families with ranges overlapping the grid cell. We additionally investigated how isolation influences diet breadth by comparing butterfly species found on islands with those on the mainland. A species was classified as an island inhabitant if its distribution was restricted to an island or group of islands, regardless of historical connection to mainland habitats. An exception were the Sunda Islands and New Guinea – several species on these islands were also present at the southern tip of the Malay Peninsula, but were considered to be island species (Gross *et al*. 2025; Kier *et al*. 2009). We used similar criteria to classify grid cells as island or mainland and characterized the proportion of species in these grid cells that were island endemics.

*Data analyses – latitudinal gradients.* To assess latitudinal trends in caterpillar diet breath, we used the Akaike Information Criterion (AIC) to compare the fit of multiple models of average diet breadth as a function of latitude across 13,316 different 100 × 100 km^2^ grid cells, namely polynomial linear models from 1 to 8 degrees and generalized additive models (GAMs). We also modeled the family-level richness of plants in the same grid cells, because past studies have shown that diet breadth often shows a negative relationship with hostplant richness.

*Data analyses – assemblage level.* For both diet dispersion and diet richness, we examined the effects of climatic, habitat, and geographic predictors in two ways: 1) using the mean diet breadths of caterpillar assemblages from 100 × 100 km^2^ grid cells as response units (n = 13,316), and 2) using individual species’ diet breadth as response units (n = 2,570). At the assemblage level, we first reduced our initial pool of 26 climatic, habitat, and host richness predictors, plus interactions between mean annual temperature (MAT) and precipitation seasonality, MAT and mean annual precipitation (MAP), and MAP and temperature seasonality, to 10 with variance inflation factors (VIFs) less than 5 using the car package in R (Fox & Weisberg 2019). Our final list included MAT (°C), mean diurnal temperature range (mean of monthly temperature range), precipitation seasonality (annual coefficient of variation), precipitation of the driest quarter (mm), percent tree cover, percent non-tree cover, net primary productivity (NPP, kg C m^-^²), elevation, family-level plant richness, and an interaction between MAT and precipitation seasonality.

We first modeled each diet breadth variable at the assemblage level in response to caterpillar species richness and phylogenetic structure (mean pairwise distance) to control for sampling effects from the global species pool and phylogenetic signal in diet breadth. We used residuals from these initial models as responses to our 10 predictor variables. For our phylogeny, we used a time-calibrated tree of 2,254 species from Kawahara *et al*. (2023) representing all families, all subfamilies and tribes, and 92% of described genera. To complete the tree, we grafted the remaining species that were not in the original tree onto the backbone topology using the function *phylo.maker* in the package V.Phylomaker2 under scenario 2 (Jin & Qian 2022), in which new tips are bound to randomly selected nodes at and below the genus- or family-level basal node. We created 100 random trees based on this method and averaged them to create a consensus tree for our assemblage-level analyses. While more explicit methods of accounting for phylogenetic signal in the analysis of assemblage and community-level trait variation exist (e.g. phylogenetic [generalized] linear mixed models, Li *et al*. 2020), the number of grid cells and species in our dataset made this approach computationally infeasible.

While caterpillar diet breadth may be directly influenced by climate or the availability of hostplants, the latter is likely influenced by climate. Therefore, we built a series of structural equation models (SEMs; Figure 2) to examine direct and indirect effects of climate and vegetation on diet breadth using the package piecewiseSEM (Lefcheck 2016). We selected our initial set of SEM predictors based on effect sizes (Cohen’s *d*) of our initial set of predictors in linear models, with a cutoff of 0.3. For diet richness, we compared nested SEMs according to AIC and multiple goodness-of-fit metrics, including χ^2^, Fisher’s C, and RMSEA to account for large sample size (Lefcheck 2016; MacCallum *et al*. 1996), testing only paths that made biological sense *a priori*. We included MAT, precipitation seasonality, their interaction, and precipitation of the driest quarter as exogenous predictor variables, and NPP, percent tree cover, and plant family richness as mediating variables. For diet dispersion, we compared nested SEMs including MAT and precipitation of the driest quarter as exogenous predictors, and vegetation variables as mediating variables. All variables were modeled in general linear models, except for plant family richness, which was modeled in a generalized linear model as a Poisson-distributed variable with a log link function.

To examine whether assemblages on islands differed in diet breadth from those in mainland habitats, we compared residual diet breadth in island grid cells (n = 445) to that in a random set of 445 mainland grid cells in a set of 1000 linear models, which we averaged based on AIC weights. Because island grid cells in our dataset are not uniformly distributed across latitudinal bands, apparent differences between island and mainland assemblages may simply reflect a broader latitudinal gradient in diet breadth, underlaid by climatic and vegetation features as described above. We thus repeated our analyses after accounting for latitudinal effects on diet breadth, using residuals from GAMs of diet breadth against latitude as our response variables.

Furthermore, diet breadth of assemblages on islands may vary depending on whether they are composed of island endemic species. Therefore, we examined the relationship between diet breadth and the proportion of species on the island that were endemic to the island or archipelago.

*Data analyses – species level.* At the species level, we modeled diet breadth as a function of climate, habitat, and plant family richness within each species’ range polygon. For each species, we overlaid butterfly range polygons onto rasters of climate and habitat data and averaged across the range, and summed the total number of plant families present within the range polygon. We narrowed our initial predictor pool down to five based on VIFs: MAT, precipitation seasonality, and their interaction, NPP, percent non-tree cover, and plant family richness. To initially account for phylogenetic signal in diet breadth, we modeled the response of species-level diet breadth to our five predictors in phylogenetic least squares models with the R package phytools (Revell 2012), using the least-squares consensus tree from our initial set of 100 random trees. Because of the greater variation in range size among butterfly species than among gid cells as replicates, we first modeled diet breadth as a function of range size in a pgls framework, and then used residuals from this model in our tests of climate and vegetation effects. Controlling for range size addressed two key factors: the tendency for large areas to host a greater diversity of plant families (Borregaard *et al*. 2016; Drakare *et al*. 2006; MacArthur & Wilson 1967), and the observation that species with broad range size are more likely to exhibit generalist feeding behaviors across their range (Hurtado *et al*. 2024; Lancaster 2020; Slatyer *et al*. 2013).

We also separately modeled the effect of island endemism on diet breadth using phylogenetic least squares. Recognizing that the full breadth of mainland ranges is much greater than that of island areas, we restricted these analyses to encompass only those mainland species whose range areas fell within 1.5 times the interquartile range of island range areas (560 island and 1,369 mainland species).

From these initial models, we derived a final set of species-level predictors to be used in SEMs of species-level diet breadth. For diet richness, we built SEMs including MAT, precipitation seasonality, and their interaction as exogenous predictors, and percent non-tree cover and plant family richness as endogenous predictors of diet richness for the full set of species, while for diet dispersion, our SEMs included MAT, precipitation seasonality and their interaction as exogenous predictors, and plant family richness as an endogenous predictor. We compared nested SEMs according to AIC, χ^2^, RMSEA, and Fisher’s C.

## Results

*Latitudinal gradients.* GAMs of diet richness (number of host families and host phylogenetic diversity) showed an asymmetric pattern around the equator, with mid-latitude peaks (26.38°S, 15.91°N and 14.54°N, respectively); dips near the equator and 47.29°N; and increases towards the poles (Fig. 1A, Fig. S1A, Table S1). The greatest change in diet richness occurred between 7.04 and 8.40°N, where phylogenetic diversity changed by 8.40 Ma latitude^-1^ and host family richness changed by 0.31 families latitude^-1^. Diet dispersion (host SES_MPD_ and host SES_MNTD_) were best explained by GAMs showing poleward increases, peaks around 6.36°N, and dips at mid-latitudes (20.93°S, 17.27°N and 22.29°S, 15.91°N, respectively; Fig. 1B, Fig. S1B, Table S1). The greatest change in diet dispersion occurred around 2.95°N, where SES_MPD_ changed by 0.011 latitude^-1^ SES_MNTD_ changed by 0.13 latitude^-1^. Modeled plant family richness showed a generally hump-shaped latitudinal pattern centered around the equator (GAM; Fig. 1, Fig S1).

**Figure 1.**
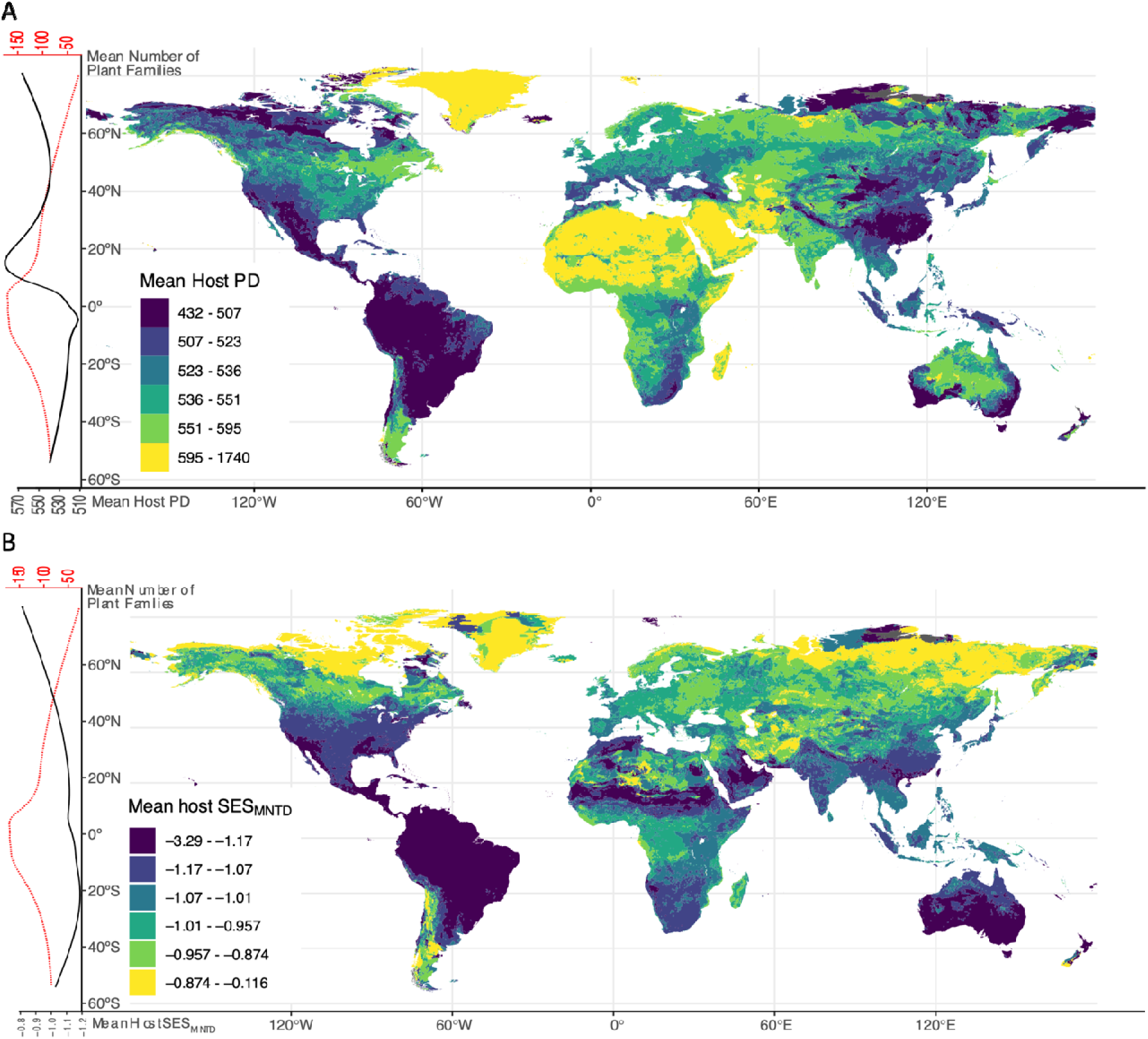
Global patterns of diet breadth in co-occurring butterfly caterpillar assemblages. Diet breadth as either mean hostplant phylogenetic diversity (PD, A) or hostplant phylogenetic dispersion (SES_MNTD_, B) is displayed globally in colors representing 6 even quantiles from the distribution of each metric. Along the left edge, we show the general latitudinal pattern of diet breadth (solid black line) overlaid with the latitudinal pattern of plant family richness (dotted red line), both represented as best fit lines from generalized additive models. Host PD showed a mid-latitude peak (14.54°N), a dip near the equator and 47.29°N, and increases towards the poles. Host SES_MNTD_ showed poleward increases, peaks around 6.36°N, and dips at mid-latitudes (22.29°S, 15.91°N). The number of host families and phylogenetic dispersion as SES_MPD_ showed similar patterns (Fig. S1)

**Figure 2.**
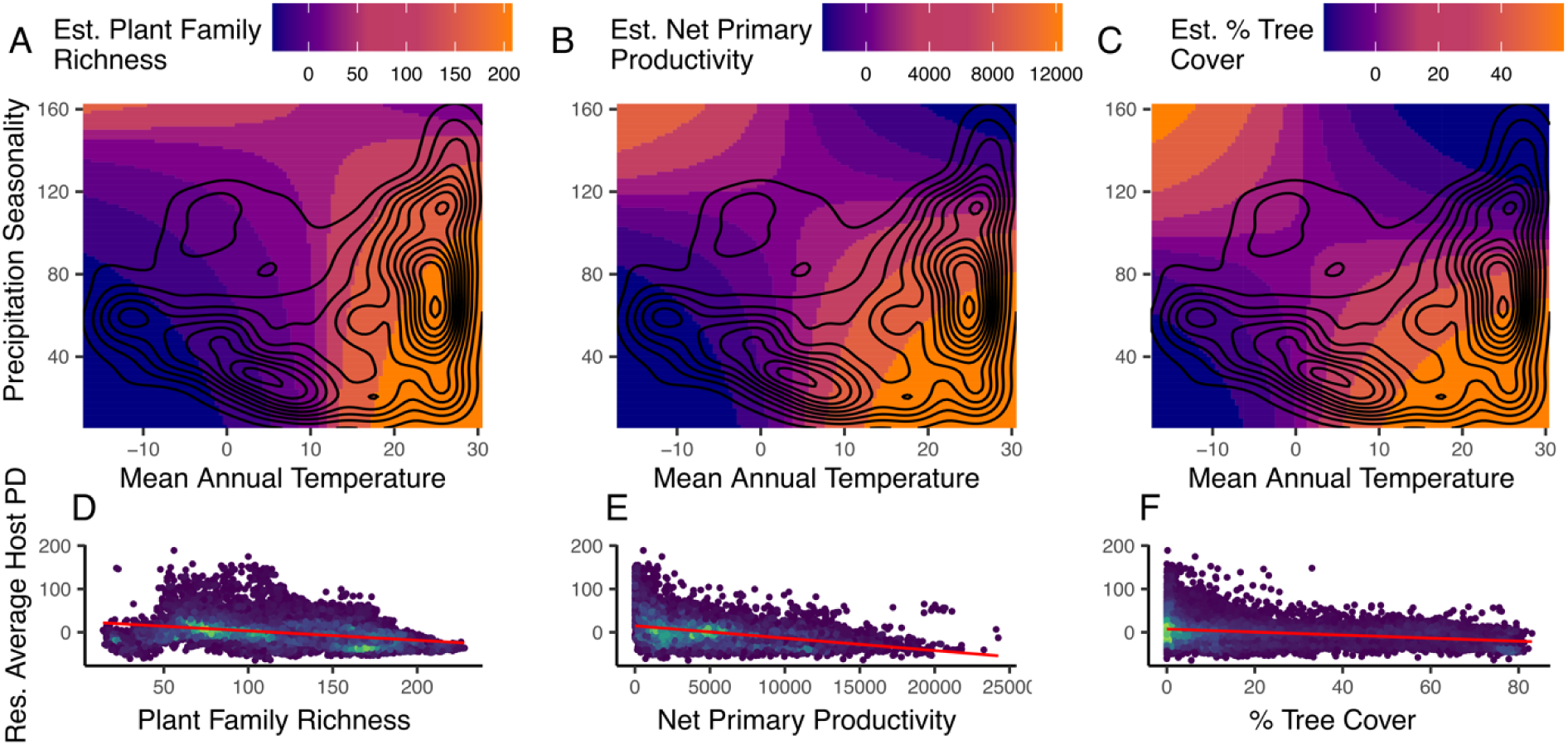
Interactions between mean annual temperature and precipitation seasonality and their effects on caterpillar diet richness. The modeled response surfaces of plant family richness (A), net primary productivity (B), and % tree cover (C) are shown as colors overlaid with the actual distribution of mean annual temperature and precipitation seasonality in 13,316 grid cells around the world (black contour lines representing point density). The observed relationships between plant family richness, net primary productivity, and % tree cover and mean caterpillar hostplant phylogenetic diversity (D-F) are shown after accounting for caterpillar species richness and phylogenetic relatedness within each grid cell. Colors in D-F represent point densities.

*Assemblage-level diet breadth.* After correcting for community phylogenetic structure and species richness, we found negative relationships between assemblage-level diet richness and local plant family richness, NPP, percent tree cover, and precipitation of the driest quarter (Table S1), with a strong interaction between precipitation seasonality and MAT (Table S2, Fig. 2). Because of our large sample sizes (n = 13,316), χ^2^ and Fisher’s C statistics indicated poor model fit. We instead focus on partial R^2^ values for diet breadth and RMSEA values (with other fit measures in Table S3), as these generally provide better indications of fit and explanatory power for the response variables of interest. Our SEMs of diet richness (phylogenetic diversity: RMSEA = 0.195, partial R^2^ = 0.33; number of hosts: RMSEA = 0.207, partial R^2^ = 0.46) showed significant direct negative effects of plant family richness and NPP on diet richness, along with a strong direct positive effect of MAT (Fig. 3A, Fig S3A). However, we also observed comparatively large indirect effects of MAT, precipitation seasonality, and their interaction mediated by plant family richness. Specifically, warm areas with low seasonal precipitation had the greatest expected NPP, richness, and tree cover, while highly seasonal warm areas and cold areas with low seasonality had the low richness, tree cover, and productivity (Fig. 2).

**Figure 3.**
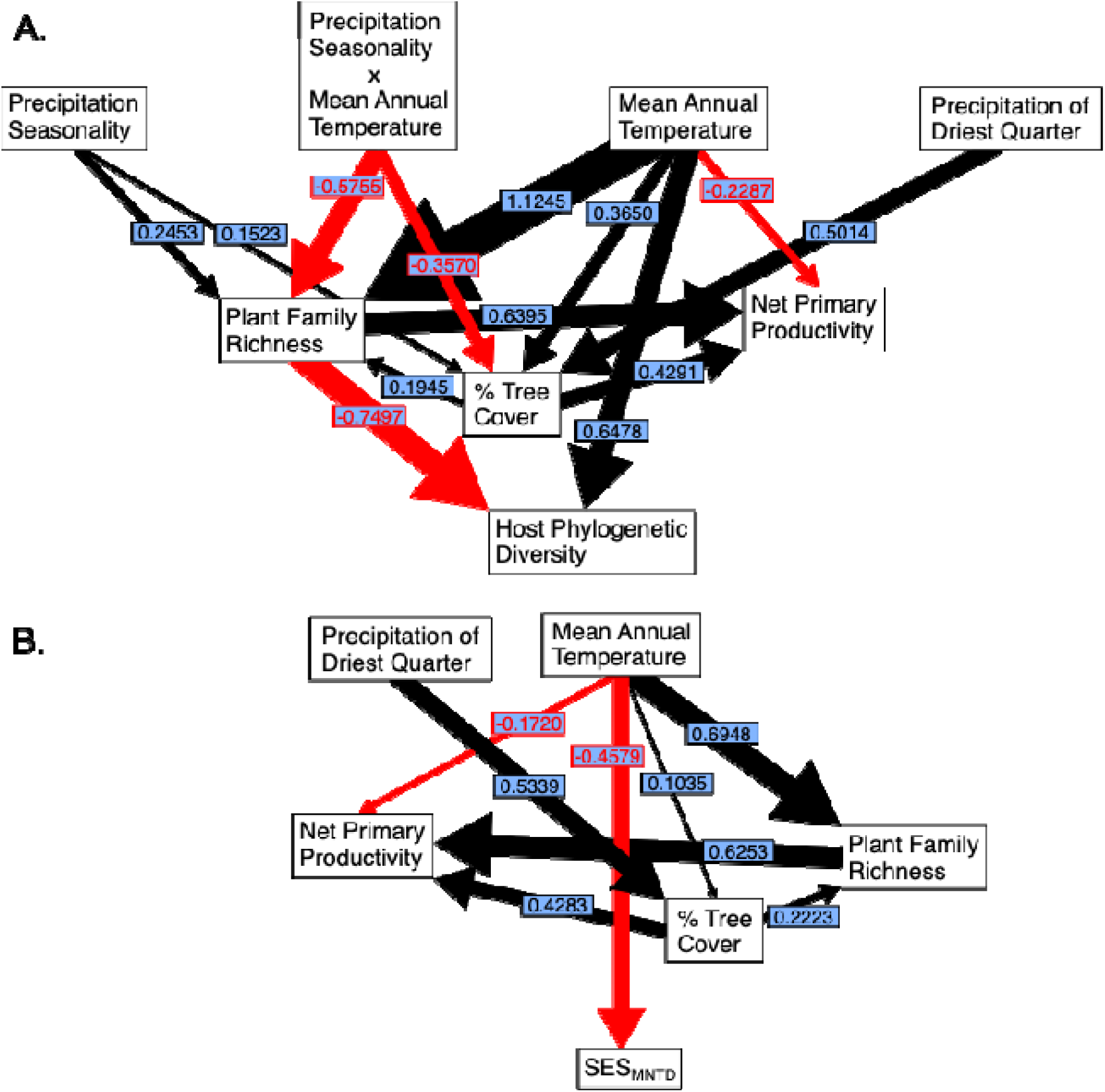
Structural equation models (SEMs) of climate and vegetation effects on assemblage- level caterpillar diet richness measured as host phylogenetic diversity (A) and diet dispersion measured as the standard effect size of host mean nearest taxon distance (B). SEMs were chosen from a set of nested identifiable models based on multiple goodness-of-fit measures (Table S3). Numbers indicate standardized path coefficients, and black arrows indicate positive effects, while red arrows indicate negative effects. For clarity, paths with significant standardized effects less than 0.1 are omitted (see Fig. S2 for all significant paths). The magnitude and direction of effects on assemblage-level diet richness quantified as the number of host families (Fig. S2A) and diet dispersion quantified as the standard effect size of host mean pairwise distance (Fig. S2C) were similar.

Our SEMs of diet dispersion performed better in terms of model fit (SES_MPD_ RMSEA = 0.0820, SES_MNTD_ RMSEA = 0.0824), but explained less variation in diet dispersion than our models of diet richness (SES_MPD_ partial R^2^ = 0.26, SES_MNTD_ partial R^2^ = 0.27; Table S3). Plant family richness and NPP had negative direct effects on diet dispersion (Fig. S2C, D). In contrast to diet richness, MAT had a negative direct effect on diet dispersion. MAT and precipitation of the driest quarter had the strongest indirect effects on diet dispersion, mediated through plant family richness and % tree cover respectively (Fig. 3B, Fig. S2C, D).

Assemblages on islands had greater diet richness on average than those in mainland cells, even after accounting for latitude (phylogenetic diversity: *R^2^* = 0.028, p < 0.001; number of hosts: *R^2^* = 0.15, p < 0.001; Fig. 4A, C), while mainland and island cells did not differ significantly in diet dispersion (SES_MPD_: *R^2^* = 0.0030, *p* = 0.136; SES_MNTD_: *R^2^* = 0.0014, *p* = 0.309). Within islands, however, we found a significant negative relationship between diet richness and island endemism – as island assemblages were more dominated by endemic species, diet richness declined (phylogenetic diversity: R^2^ = 0.077, p < 0.001; number of hosts: R^2^ = 0.027, p < 0.001; Fig. 4B, D), even after accounting for latitude.

**Figure 4.**
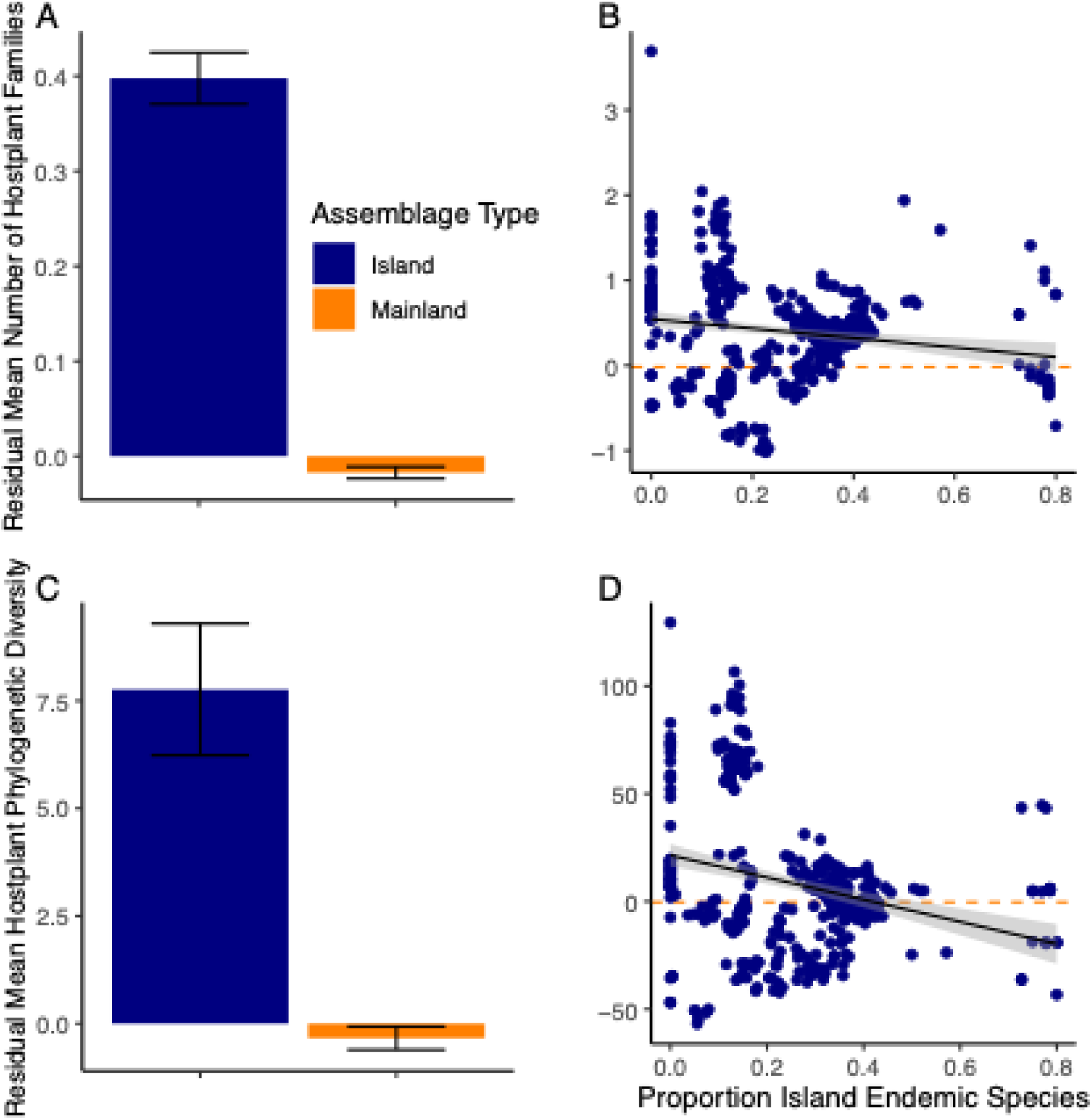
Diet richness of island and mainland caterpillar assemblages. After accounting for caterpillar species richness, phylogenetic structure, and latitude, caterpillar assemblages on islands on average fed on more plant families (A; R^2^ = 0.15, p < 0.001) and more phylogenetically diverse plant families (C; R^2^ = 0.028, p < 0.001) than those in mainland areas. Across the 445 island assemblages in our dataset, those with more island endemic species on average fed on fewer (B; R^2^ = 0.027, p < 0.001) and less phylogenetically diverse (D; R^2^ = 0.077, p < 0.001) plant families. In panels A and C, we show the mean and standard error of diet richness for all mainland and island assemblages, but we iteratively tested for an island effect using a comparable set of 445 randomly chosen mainland assemblages. In panels B and D, the dashed horizontal lines represent the mean residual number of hosts and phylogenetic diversity of mainland assemblages, respectively.

*Species-level diet breadth.* For species-level diet richness, we found significant positive interactions between MAT and precipitation seasonality (number of families p < 0.001, host PD p < 0.001), and positive correlations with plant family richness (number of families p < 0.001, host PD p < 0.001) even after accounting for variation in range size. Species-level diet dispersion was best explained by plant family richness (SES_MPD_ p < 0.001, SES_MNTD_ p = 0.00752) and the interaction between MAT and precipitation seasonality (SES_MPD_ p < 0.001, SES_MNTD_ p = 0.00365; Fig. S3). For the subset of island and mainland species with comparable ranges, island species did not differ significantly in diet richness or diet dispersion.

Our SEMs did not explain much of the variation in diet breadth (number of hosts partial R^2^ = 0.03; host PD partial R^2^ = 0.03; SES_MPD_ partial R^2^ < 0.01; SES_MPD_ partial R^2^ < 0.01).

Within-range hostplant family richness was the only consistent direct predictor of diet breadth across metrics, although we also recovered direct effects of MAT on diet richness. The number of hostplants was more strongly directly influenced by plant family richness, while host PD was more strongly directly influenced by MAT (Fig. S3).

## Discussion

*Latitudinal gradients in diet breadth.* We observed a peak in generalist assemblages around 15°N for both metrics of diet richness (Fig. 1A, Fig. S1A) rather than a monotonic latitudinal gradient in diet richness across caterpillar assemblages, with generalist assemblages at high latitudes grading into specialists at the equator. The strongest climatic predictors of diet breadth were precipitation seasonality and an interaction between precipitation seasonality and mean annual temperature (MAT; Table S1, Fig. 3A). Our 15°N peak in generalist caterpillars corresponds to western Mexico, the Indian subcontinent, and the Sahel, all of which experience high interannual variation in precipitation (a coefficient of variation up to 173.17 in Sudan).

Accordingly, we observed a generalism “hotspot” in northern Africa that roughly corresponds to this region (Fig. 1A, Fig. S1A). *Climate effects on diet breadth.* We observed a direct positive effect of MAT on diet richness (Fig. 3A, Fig. S2A), but no direct effect of climate seasonality, despite high generalism where precipitation seasonality is highest. The MAT effect could conceivably be the result of an initially hypothesized mechanism – that more intense seasonal fluctuations in climate would select for generalism by limiting the flight time of ovipositing females – except that MAT is negatively correlated with temperature seasonality (Fick & Hijmans 2017). Instead, extreme heat, especially in areas with low tree cover (such as the Sahara and Sahel, where we observed the most generalist assemblages) may similarly limit flight time and female hostplant choice (Neu *et al*. 2021). Alternatively, our results may reflect greater plasticity and intraspecific variation in hostplant use in hotter climates. Numerous studies have demonstrated that butterflies and other insects are able to alter their choice of hostplant in extreme temperatures in the interest of increasing heat tolerance, accommodating physiological changes associated with increased temperatures, or accommodating differences among hostplants in heat tolerance (Bodlah *et al*. 2016; Clissold *et al*. 2013; Hellmann 2002).

Diet dispersion exhibited latitudinal patterns distinct from diet richness (Fig. 1B, Fig. S1B), and was driven by a direct negative effect of MAT that was equivalent to (SES_MPD_; Fig. S2C) or greater than (SES_MNTD_; Fig. 3B) that of plant family richness. While assemblage-level average dispersion was consistently negative (clustered), areas with greater MAT were more negative. Here, MAT may act more strongly as a driver of increased specialization by directly increasing rates of evolution and diversification (Skeels *et al*. 2023), allowing caterpillars to specialize on narrow regions of the plant phylogeny in a pattern that is not simply summarized by quantifying branch lengths or numbers of taxa. Alternatively, the direct effect of MAT may reflect its role as an ecological filter that restricts the phylogenetic dispersion of available hostplant families that is realized as caterpillar diet dispersion. For example, if heat tolerance is phylogenetically conserved, high temperatures may restrict the subset of families from a regional pool that are able to persist within a grid cell, and these families would be more closely related than expected from a random draw from the regional pool (Lebrija-Trejos *et al*. 2010). The fact that we did not observe similar direct effects of MAT on diet dispersion at the species level but did for diet richness (Fig. S3) may support this plant assemblage-level mechanism of specialization.

*Plant diversity effects on diet breadth.* While precipitation seasonality was the strongest individual climatic predictor of diet richness, it did not directly exert any strong influence on diet richness (Fig. 3A, Fig. S2A). Instead, precipitation seasonality interacted with MAT to affect plant family richness, which was the strongest direct driver of diet richness (Fig. 2, Fig. 3A, Fig. S2A). When precipitation seasonality was high, plant family richness was moderate across all temperature regimes. However, when precipitation seasonality was low, plant family richness was highest at high temperatures, reflecting the consistently warm and wet climate of the equatorial tropics, where we observed the maximum average plant family richness (Fig. 1, Fig. S1) and the minimum average diet richness (Fig. 1A, Fig. S1A).

In areas of high plant diversity, specialization is thought to be favored for several reasons. First, high plant diversity allows herbivores to be more selective of their hostplants, favoring specialization in the interest of reducing competition for food and oviposition sites at high densities (Benson 1978; Bird *et al*. 2019; Shapiro & Cardé 1970). High herbivore density may itself be a possible function of high primary productivity (Kaspari *et al*. 2000; Storch *et al*. 2018), which also exerted small but significant negative effects on diet richness (Fig. S2A, B). Higher-intensity trophic interactions in diverse tropical habitats can select for strong and distinct plant chemical defenses (Endara *et al*. 2023; Sun *et al*. 2024), in turn selecting for specialist herbivores uniquely equipped to metabolize and/or sequester those defenses (Ehrlich & Raven 1964; van der Linden *et al*. 2021), while metabolic costs of host-switching tend to favor narrow diets (Braga *et al*. 2021; Janz & Nylin 1998; Nishida 2002).

*Island endemism drives specialization.* Caterpillar assemblages on islands appeared to be much less specialized on their hostplants than those in mainland areas on average (Fig. 4A, 4C), consistent with the hypothesis that most assemblages on islands that we observed consisted of species with broader ranges that include islands as well as mainland regions. Butterfly populations at range edges often feed on distinctive or novel hostplants, expanding the species-average niche breadth (Lancaster 2020). Because our compiled hostplant dataset is aggregated across multiple populations, expansion into a wider geographic area equates to broader niches on average across the range of the entire species, despite narrow diet breadth that may occur at range edges (Hurtado *et al*. 2024; Lancaster 2020). For individual grid cells containing species with wide inter-population variation in hostplants, generalism is the norm.

However, assemblages with a larger proportion of island-endemic species tended to be more specialized on average (Fig. 4B, 4D), hinting that island endemism plays a significant role in driving assemblage-level specialization. Island endemics may drive specialization due to longer histories of isolation from mainland habitats. Non-endemic island species likely represent recent colonization events in marginal habitats; they may be specialized at the population level within their broader range (Lancaster 2020), but they have had insufficient time to become reproductively isolated as distinctive “neo-endemic” species (Gillespie & Roderick 2002; Losos & Ricklefs 2009). Furthermore, the geographic isolation and small size of islands with high endemism contribute to limited availability of hostplants (Borregaard *et al*. 2016; MacArthur & Wilson 1967), which may also drive specialization. Limited hostplant availability is provided as one explanation for why there are multiple examples of carnivory in Lepidoptera on islands (Montgomery 1982; Rubinoff & Haines 2005), while it is comparatively much rarer on mainlands. Species endemic to larger islands with historic connections to continents likely represent “paleo-endemic” lineages, with origins that predate isolation, and the mechanisms that drive specialization in these regions are less clear.

While our results on the global pattern of diet breadth differs from that of a prior study (Forister *et al*. 2015), the underlying mechanisms are consistent: global patterns in plant diversity play a key role in shaping caterpillar diet breadth, and climate has a significant indirect and direct impact. The roles of precipitation seasonality and temperature are testament both to the importance of climate in dictating the evolution and ecology of terrestrial ectotherms (Archibald *et al*. 2010; Chowdhury *et al*. 2021; Sunday *et al*. 2010), and to the vulnerability of butterfly and other insect communities to global change as these climate regimes shift (Minev-Benzecry & Daru 2024; Wang *et al*. 2024). Similarly, the influence of endemism and range size on diet breadth of island caterpillars emphasize the potential impact of range expansions and species invasions on insect communities and food web dynamics in these isolated habitats (Lozan *et al*. 2008; New 2008). While our results represent a broad-scale exploration of patterns and processes that have their roots in the deeper evolutionary histories of butterflies and plants, we anticipate that our work will provide valuable insights into how these processes shape butterfly and hostplant responses to global change.

---

**Figure S1.**
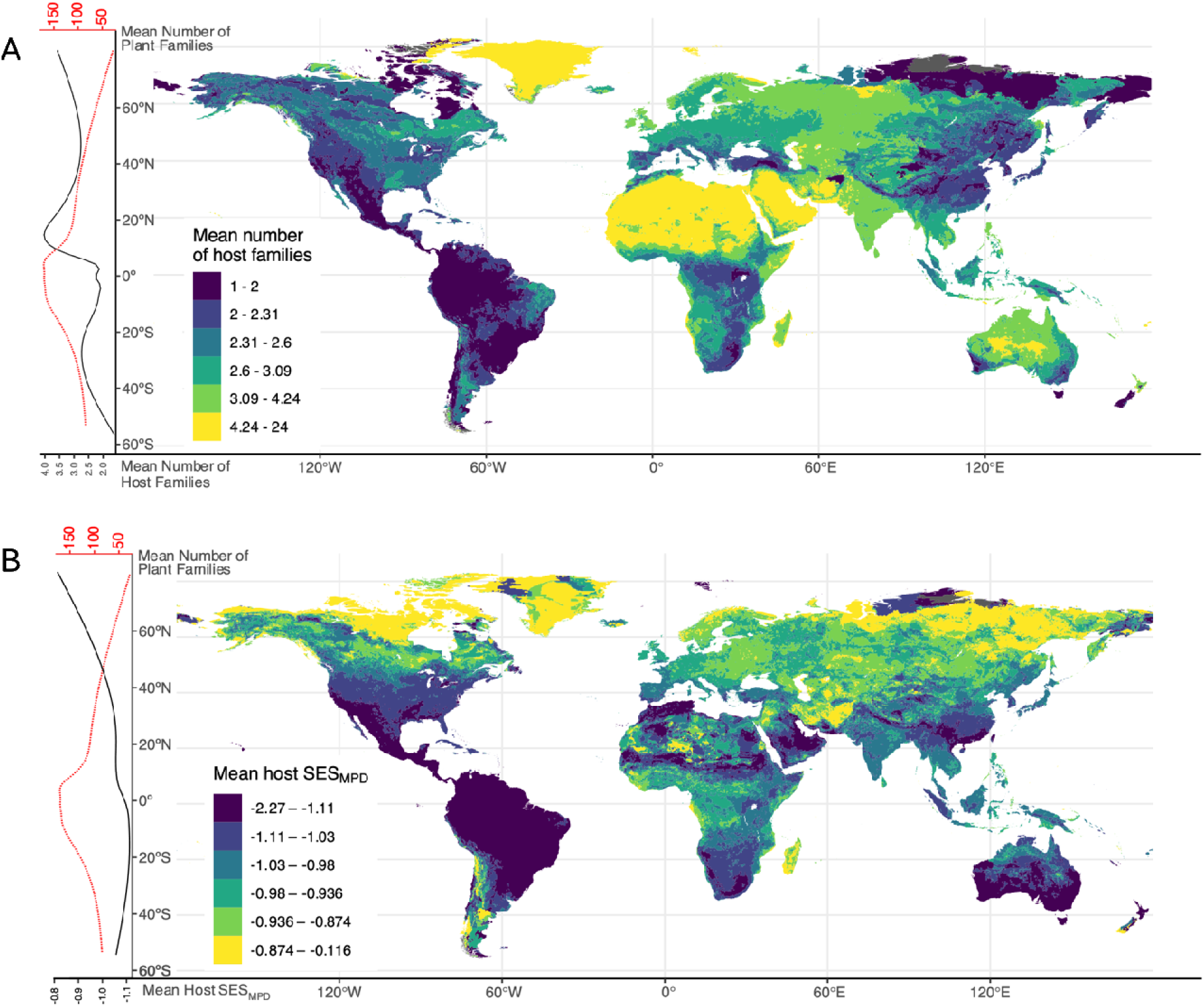
Global patterns of diet breadth in co-occurring butterfly caterpillar assemblages. Diet breadth as either the mean number of hostplant families (A) or hostplant phylogenetic dispersion (SES_MPD_, B) is displayed globally in colors representing 6 even quantiles from the distribution of each metric. Along the left edge, we show the general latitudinal pattern of diet breadth (solid black line) overlaid with the latitudinal pattern of plant family richness (dotted red line), both represented as best fit lines from generalized additive models. Number of host families showed a mid-latitude peak (15.90°N), a dip near the equator, and increases towards the poles. Host SES_MPD_ showed poleward increases, peaks around 6.36°N, and a mid-latitude dip (20.93°S). Host phylogenetic diversity and phylogenetic dispersion as SES_MNTD_ showed similar patterns (Fig. 1)

**Figure S2.**
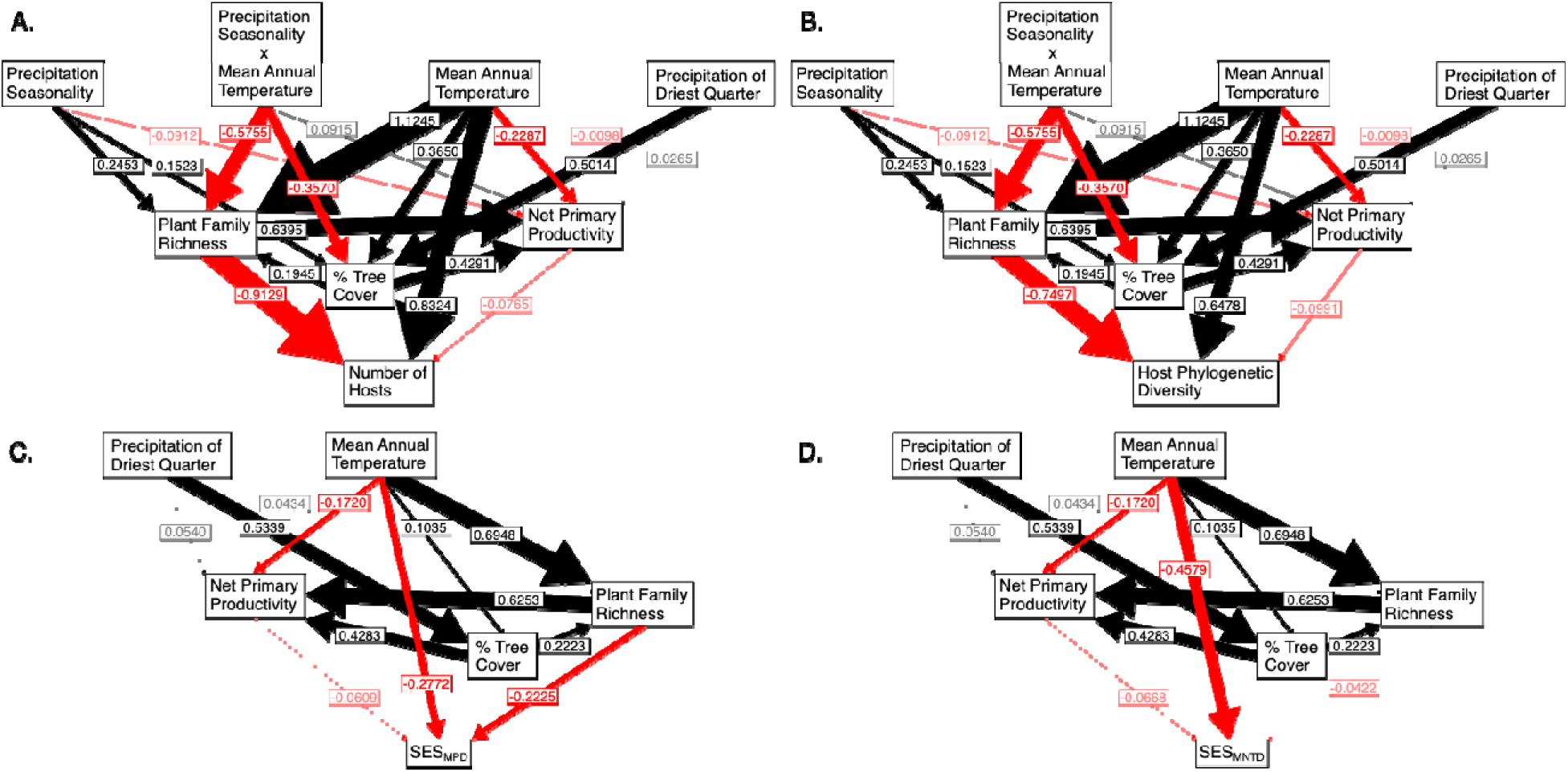
Structural equation models (SEMs) of climate and vegetation effects on assemblage- level caterpillar diet richness measured as host family richness (A) and host phylogenetic diversity (B); and diet dispersion measured as the standard effect size of host mean pairwise distance (C) and host mean nearest taxon distance (D). SEMs were chosen from a set of nested identifiable models based on multiple goodness-of-fit measures (Table S3). All significant paths from the selected model are shown here; numbers indicate standardized path coefficients, and black arrows indicate positive effects, while red arrows indicate negative effects. For clarity, paths with standardized effects less than 0.1 are shown with lowered opacity.

**Figure S3.**
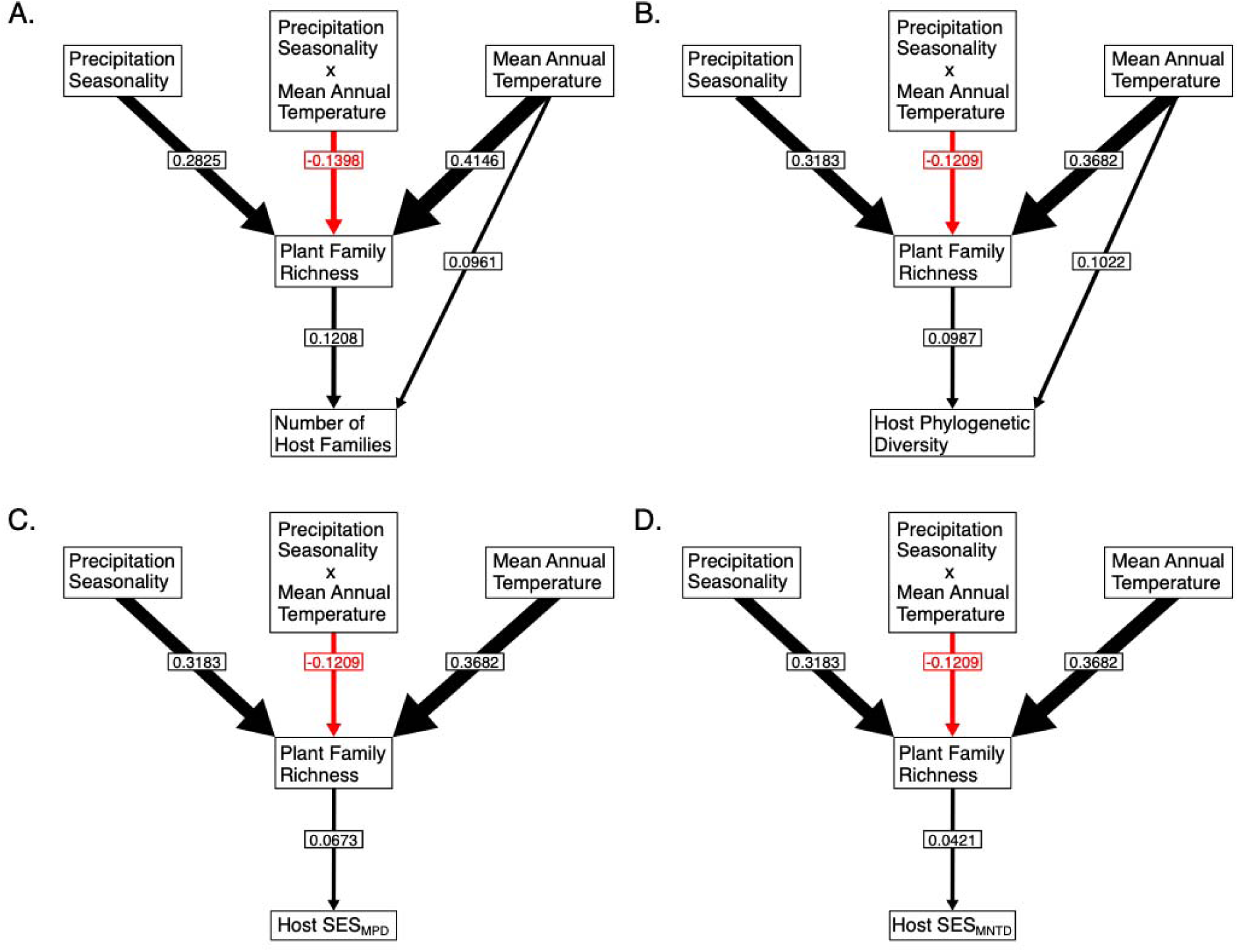
Structural equation models (SEMs) of climate and vegetation effects on species-level caterpillar diet richness measured as host family richness (A) and host phylogenetic diversity (B), and diet dispersion measured as the standard effect size of host mean pairwise phylogenetic distance (C) and mean nearest taxon difference (D). SEMs were chosen from a set of nested identifiable models based on multiple goodness-of-fit measures (Table S4).

**Table S1.**
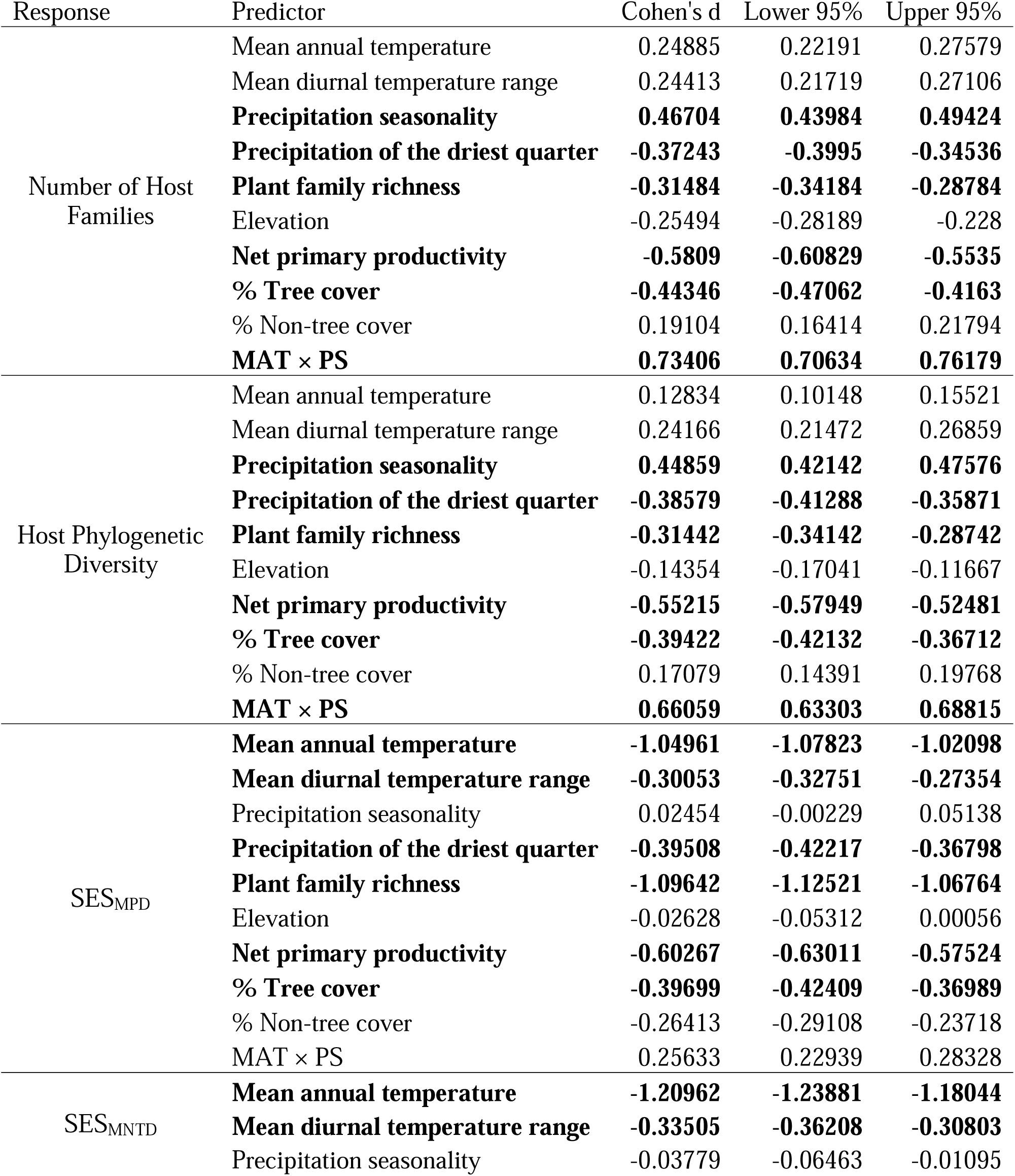
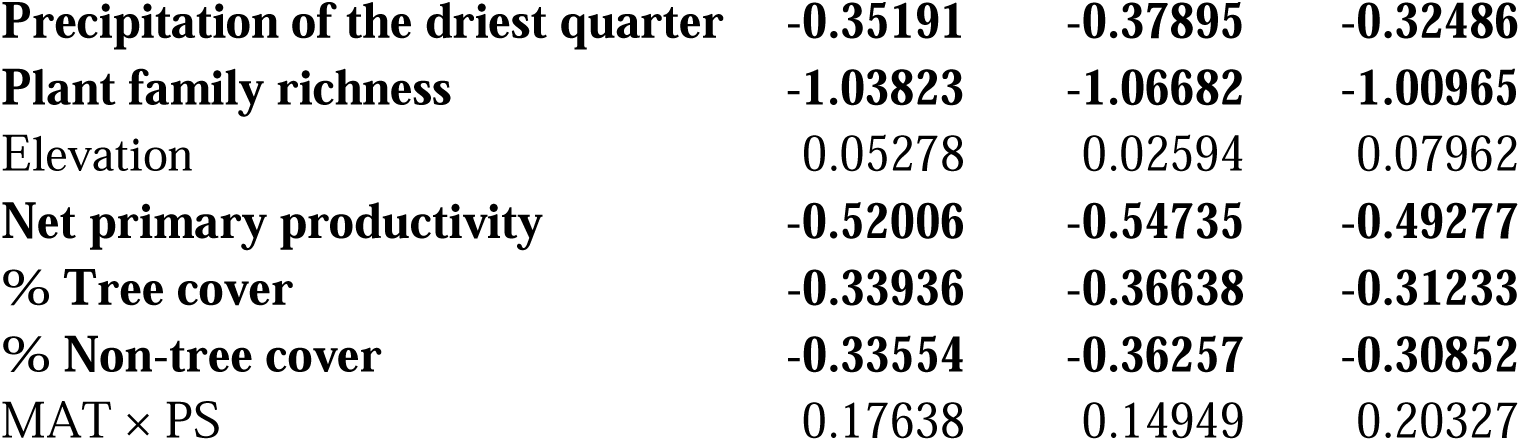
Standardized effect sizes (Cohen’s *d*) of climate and vegetation predictors of caterpillar diet breadth in linear models. Effect sizes with an absolute value greater than 0.3 (bolded) were incorporated into structural equation models.

| Response | Predictor | Cohen's d | Lower 95% | Upper 95% |
| --- | --- | --- | --- | --- |
| Number of Host Families | Mean annual temperature | 0.24885 | 0.22191 | 0.27579 |
|  | Mean diurnal temperature range | 0.24413 | 0.21719 | 0.27106 |
|  | **Precipitation seasonality** | **0.46704** | **0.43984** | **0.49424** |
|  | **Precipitation of the driest quarter** | **-0.37243** | **-0.3995** | **-0.34536** |
|  | **Plant family richness** | **-0.31484** | **-0.34184** | **-0.28784** |
|  | Elevation | -0.25494 | -0.28189 | -0.228 |
|  | **Net primary productivity** | **-0.5809** | **-0.60829** | **-0.5535** |
|  | **% Tree cover** | **-0.44346** | **-0.47062** | **-0.4163** |
|  | % Non-tree cover | 0.19104 | 0.16414 | 0.21794 |
|  | **MAT × PS** | **0.73406** | **0.70634** | **0.76179** |
| Host Phylogenetic Diversity | Mean annual temperature | 0.12834 | 0.10148 | 0.15521 |
|  | Mean diurnal temperature range | 0.24166 | 0.21472 | 0.26859 |
|  | **Precipitation seasonality** | **0.44859** | **0.42142** | **0.47576** |
|  | **Precipitation of the driest quarter** | **-0.38579** | **-0.41288** | **-0.35871** |
|  | **Plant family richness** | **-0.31442** | **-0.34142** | **-0.28742** |
|  | Elevation | -0.14354 | -0.17041 | -0.11667 |
|  | **Net primary productivity** | **-0.55215** | **-0.57949** | **-0.52481** |
|  | **% Tree cover** | **-0.39422** | **-0.42132** | **-0.36712** |
|  | % Non-tree cover | 0.17079 | 0.14391 | 0.19768 |
|  | **MAT × PS** | **0.66059** | **0.63303** | **0.68815** |
| SES<sub>MPD</sub> | **Mean annual temperature** | **-1.04961** | **-1.07823** | **-1.02098** |
|  | **Mean diurnal temperature range** | **-0.30053** | **-0.32751** | **-0.27354** |
|  | Precipitation seasonality | 0.02454 | -0.00229 | 0.05138 |
|  | **Precipitation of the driest quarter** | **-0.39508** | **-0.42217** | **-0.36798** |
|  | **Plant family richness** | **-1.09642** | **-1.12521** | **-1.06764** |
|  | Elevation | -0.02628 | -0.05312 | 0.00056 |
|  | **Net primary productivity** | **-0.60267** | **-0.63011** | **-0.57524** |
|  | **% Tree cover** | **-0.39699** | **-0.42409** | **-0.36989** |
|  | % Non-tree cover | -0.26413 | -0.29108 | -0.23718 |
|  | MAT × PS | 0.25633 | 0.22939 | 0.28328 |
| SES<sub>MNTD</sub> | **Mean annual temperature** | **-1.20962** | **-1.23881** | **-1.18044** |
|  | **Mean diurnal temperature range** | **-0.33505** | **-0.36208** | **-0.30803** |
|  | Precipitation seasonality | -0.03779 | -0.06463 | -0.01095 |

|  |  |  |  |
| --- | --- | --- | --- |
| **Precipitation of the driest quarter** | **-0.35191** | **-0.37895** | **-0.32486** |
| **Plant family richness** | **-1.03823** | **-1.06682** | **-1.00965** |
| Elevation | 0.05278 | 0.02594 | 0.07962 |
| **Net primary productivity** | **-0.52006** | **-0.54735** | **-0.49277** |
| **% Tree cover** | **-0.33936** | **-0.36638** | **-0.31233** |
| **% Non-tree cover** | **-0.33554** | **-0.36257** | **-0.30852** |
| MAT × PS | 0.17638 | 0.14949 | 0.20327 |

**Table S2.** Latitudinal optima of caterpillar diet breadth – measured as number of host families, host phylogenetic diversity, and host phylogenetic dispersion (SES_MPD_ and SES_MNTD_) – and plant family richness, from generalized additive models.

| Response | Local Optimum | Predicted value |
| --- | --- | --- |
| Number of Host Families | 47.286779°N | 2.761788 |
|  | 15.908399°N | 4.040527 |
|  | 3.629902°N | 2.143608 |
|  | 0.901347°N | 2.264598 |
|  | 1.827208°S | 2.092453 |
|  | 26.384201°S | 2.74046 |
| Host Phylogenetic Diversity | 47.286779°N | 538.6848 |
|  | 14.544121°N | 583.5817 |
|  | 4.555763°S | 511.5399 |
| $SES_{MPD}$ | 17.272676°N | -1.065329 |
|  | 6.358457°N | -1.052092 |
|  | 5.920040°S | -1.110561 |
|  | 11.377150°S | -1.106413 |
|  | 20.927092°S | -1.110569 |
| $SES_{MNTD}$ | 15.908399°N | -1.131788 |
|  | 6.358457°N | -1.094601 |
|  | 5.920040°S | -1.16042 |
|  | 8.648595°S | -1.155973 |
|  | 22.291369°S | -1.187228 |
| Plant Family Richness | 3.629902°N | 171.1766 |
|  | 0.901347°N | 168.8615 |
|  | 1.827208°S | 170.8485 |

## References

1. Ackermann, M. & Doebeli, M. (2004). Evolution of Niche Width and Adaptive Diversification. Evolution, 58, 2599–2612.

2. Agosta, S.J., Janz, N. & Brooks, D.R. (2010). How specialists can be generalists: resolving the “parasite paradox” and implications for emerging infectious disease. Zoologia (Curitiba), 27, 151–162.

3. Agrawal, A.A. & Hastings, A.P. (2023). Tissue-specific plant toxins and adaptation in a specialist root herbivore. Proceedings of the National Academy of Sciences, 120, e2302251120.

4. Archibald, S.B., Bossert, W.H., Greenwood, D.R. & Farrell, B.D. (2010). Seasonality, the latitudinal gradient of diversity, and Eocene insects. Paleobiology, 36, 374–398.

5. Augustine, K.E. & Kingsolver, J.G. (2018). Biogeography and phenology of oviposition preference and larval performance of Pieris virginiensis butterflies on native and invasive host plants. Biol Invasions, 20, 413–422.

6. Barnes, E.E. & Murphy, S.M. (2023). Bottom-up and top-down pressures mediate competition between two generalist insects. Ecology, 104, e3957.

7. Becker, L., Blüthgen, N. & Drossel, B. (2022). Stochasticity Leads to Coexistence of Generalists and Specialists in Assembling Mutualistic Communities. The American Naturalist, 200, 303–315.

8. Benson, W.W. (1978). RESOURCE PARTITIONING IN PASSION VINE BUTTERFLIES. Evolution, 32, 493–518.

9. Bird, G., Kaczvinsky, C., Wilson, A.E. & Hardy, N.B. (2019). When do herbivorous insects compete? A phylogenetic meta-analysis. Ecology Letters, 22, 875–883.

10. Bodlah, M.A., Zhu, A.-X. & Liu, X.-D. (2016). Host choice, settling and folding leaf behaviors of the larval rice leaf folder under heat stress. Bulletin of Entomological Research, 106, 809–817.

11. Borregaard, M.K., Matthews, T.J. & Whittaker, R.J. (2016). The general dynamic model: towards a unified theory of island biogeography? Global Ecology and Biogeography, 25, 805–816.

12. Braga, M.P., Janz, N., Nylin, S., Ronquist, F. & Landis, M.J. (2021). Phylogenetic reconstruction of ancestral ecological networks through time for pierid butterflies and their host plants. Ecology Letters, 24, 2134–2145.

13. Braga, M.P., Landis, M.J., Nylin, S., Janz, N. & Ronquist, F. (2020). Bayesian Inference of Ancestral Host–Parasite Interactions under a Phylogenetic Model of Host Repertoire Evolution. Systematic Biology, 69, 1149–1162.

14. Bruijning, M., Metcalf, C.J.E. & Visser, M.D. (2024). Closing the gap in the Janzen–Connell hypothesis: What determines pathogen diversity? Ecology Letters, 27, e14316.

15. Cai, L., Kreft, H., Taylor, A., Denelle, P., Schrader, J., Essl, F., et al. (2023). Global models and predictions of plant diversity based on advanced machine learning techniques. New Phytologist, 237, 1432–1445.

16. Chowdhury, S., Fuller, R.A., Dingle, H., Chapman, J.W. & Zalucki, M.P. (2021). Migration in butterflies: a global overview. Biological Reviews, 96, 1462–1483.

17. Clissold, F.J., Coggan, N. & Simpson, S.J. (2013). Insect herbivores can choose microclimates to achieve nutritional homeostasis. Journal of Experimental Biology, 216, 2089–2096.

18. Daru, B.H. (2024a). A global database of butterfly species native distributions. *Ecology*, n/a, e4462.

19. Daru, B.H. (2024b). Predicting undetected native vascular plant diversity at a global scale. Proceedings of the National Academy of Sciences, 121, e2319989121.

20. Daru, B.H., Karunarathne, P. & Schliep, K. (2020). phyloregion: R package for biogeographical regionalization and macroecology. Methods in Ecology and Evolution, 11, 1483–1491.

21. Day, E.H., Hua, X. & Bromham, L. (2016). Is specialization an evolutionary dead end? Testing for differences in speciation, extinction and trait transition rates across diverse phylogenies of specialists and generalists. Journal of Evolutionary Biology, 29, 1257– 1267.

22. Devictor, V., Julliard, R. & Jiguet, F. (2008). Distribution of specialist and generalist species along spatial gradients of habitat disturbance and fragmentation. Oikos, 117, 507–514.

23. Drakare, S., Lennon, J.J. & Hillebrand, H. (2006). The imprint of the geographical, evolutionary and ecological context on species–area relationships. Ecology Letters, 9, 215–227.

24. Dubayah, R.O., Luthcke, S.B., Sabaka, T.J., Nicholas, J.B., Preaux, S. & Hofton, M.A. (2021) GEDI L3 Gridded Land Surface Metrics, Version 2.

26. Dyer, L.A. & Forister, M.L. (2019). Challenges and advances in the study of latitudinal gradients in multitrophic interactions, with a focus on consumer specialization. *Current Opinion in Insect Science*, Ecology • Parasites/Parasitoids/Biological control, 32, 68–76.

27. Ehrlich, P.R. & Raven, P.H. (1964). Butterflies and Plants: A Study in Coevolution. Evolution, 18, 586–608.

28. Endara, M.-J., Forrister, D.L. & Coley, P.D. (2023). The Evolutionary Ecology of Plant Chemical Defenses: From Molecules to Communities. *Annual Review of Ecology*, Evolution, and Systematics, 54, 107–127.

29. Faith, D.P. (1992). Conservation evaluation and phylogenetic diversity. Biological Conservation, 61, 1–10.

30. Fick, S.E. & Hijmans, R.J. (2017). WorldClim 2: new 1-km spatial resolution climate surfaces for global land areas. International Journal of Climatology, 37, 4302–4315.

31. Fordyce, J.A. (2010). Host shifts and evolutionary radiations of butterflies. Proc Biol Sci, 277, 3735–3743.

32. Forister, M.L., Novotny, V., Panorska, A.K., Baje, L., Basset, Y., Butterill, P.T., et al. (2015). The global distribution of diet breadth in insect herbivores. Proceedings of the National Academy of Sciences, 112, 442–447.

33. Fox, J. & Weisberg, S. (2019). An R Companion to Applied Regression. 3rd edn. Sage, Thousand Oaks, CA. *GBIF*: Global Biogeographic Information Facility. (2023). . Available at: https://www.gbif.org/. Last accessed 23 February 2023.

34. Gillespie, R.G. & Roderick, G.K. (2002). Arthropods on Islands: Colonization, Speciation, and Conservation. Annual Review of Entomology, 47, 595–632.

35. Gross, C.P., Wright, A.M. & Daru, B.H. (2025). A global biogeographic regionalization for butterflies. Philosophical Transactions of the Royal Society B: Biological Sciences, 380, 20230211.

36. Hardy, N.B. & Otto, S.P. (2014). Specialization and generalization in the diversification of phytophagous insects: tests of the musical chairs and oscillation hypotheses. Proceedings of the Royal Society B: Biological Sciences, 281, 20132960.

37. Hargreaves, A.L. (2024). Geographic Gradients in Species Interactions: From Latitudinal Patterns to Ecological Mechanisms. Annual Review of Ecology, Evolution, and Systematics, 55, 369–393.

38. Hellmann, J.J. (2002). The effect of an environmental change on mobile butterfly larvae and the nutritional quality of their hosts. Journal of Animal Ecology, 71, 925–936.

39. Hembry, D.H., Bennett, G., Bess, E., Cooper, I., Jordan, S., Liebherr, J., et al. (2021). Insect Radiations on Islands: Biogeographic Pattern and Evolutionary Process in Hawaiian Insects. The Quarterly Review of Biology, 96, 247–296.

40. Hurtado, P., Aragón, G., Vicente, M., Dalsgaard, B., Krasnov, B.R. & Calatayud, J. (2024). Generalism in species interactions is more the consequence than the cause of ecological success. Nat Ecol Evol, 8, 1602–1611.

41. Janz, N. & Nylin, S. (1998). BUTTERFLIES AND PLANTS: A PHYLOGENETIC STUDY. Evolution, 52, 486–502.

42. Jin, Y. & Qian, H. (2022). V.PhyloMaker2: An updated and enlarged R package that can generate very large phylogenies for vascular plants. Plant Diversity, 44, 335–339.

43. Kaspari, M., Alonso, L. & O’Donnell, S. (2000). Three energy variables predict ant abundance at a geographical scale. Proceedings of the Royal Society of London. Series B: Biological Sciences, 267, 485–489.

44. Kawahara, A.Y., Storer, C., Carvalho, A.P.S., Plotkin, D.M., Condamine, F.L., Braga, M.P., et al. (2023). A global phylogeny of butterflies reveals their evolutionary history, ancestral hosts and biogeographic origins. Nat Ecol Evol, 7, 903–913.

45. Kembel, S.W., Cowan, P.D., Helmus, M.R., Cornwell, W.K., Morlon, H., Ackerly, D.D., et al. (2010). Picante: R tools for integrating phylogenies and ecology. Bioinformatics, 26, 1463–1464.

46. Kier, G., Kreft, H., Lee, T.M., Jetz, W., Ibisch, P.L., Nowicki, C., et al. (2009). A global assessment of endemism and species richness across island and mainland regions. Proceedings of the National Academy of Sciences, 106, 9322–9327.

47. Lancaster, L.T. (2020). Host use diversification during range shifts shapes global variation in Lepidopteran dietary breadth. Nat Ecol Evol, 4, 963–969.

48. Lebrija-Trejos, E., Pérez-García, E.A., Meave, J.A., Bongers, F. & Poorter, L. (2010). Functional traits and environmental filtering drive community assembly in a species-rich tropical system. Ecology, 91, 386–398.

49. Lefcheck, J.S. (2016). piecewiseSEM: Piecewise structural equation modelling in r for ecology, evolution, and systematics. Methods in Ecology and Evolution, 7, 573–579.

50. Li, D., Dinnage, R., Nell, L.A., Helmus, M.R. & Ives, A.R. (2020). phyr: An r package for phylogenetic species-distribution modelling in ecological communities. Methods in Ecology and Evolution, 11, 1455–1463.

51. van der Linden, C.F.H., WallisDeVries, M.F. & Simon, S. (2021). Great chemistry between us: The link between plant chemical defenses and butterfly evolution. Ecology and Evolution, 11, 8595–8613.

52. Losos, J.B. & Ricklefs, R.E. (2009). Adaptation and diversification on islands. Nature, 457, 830– 836.

53. Louca, S. (2021). Phylogeographic Estimation and Simulation of Global Diffusive Dispersal. Syst Biol, 70, 340–359.

54. Lozan, A.I., Monaghan, M.T., Spitzer, K., Jaroš, J., Žurovcová, M. & Brož, V. (2008). DNA- based confirmation that the parasitic wasp Cotesia glomerata (Braconidae, Hymenoptera) is a new threat to endemic butterflies of the Canary Islands. Conserv Genet, 9, 1431– 1437.

55. MacArthur, R.H. & Wilson, E.O. (1967). The Theory of Island Biogeography. Princeton Monographs in Population Biology. 1st edn. Princeton University Press, Princeton, NJ.

56. MacCallum, R.C., Browne, M.W. & Sugawara, H.M. (1996). Power analysis and determination of sample size for covariance structure modeling. Psychological Methods, 1, 130–149.

57. Minev-Benzecry, S. & Daru, B.H. (2024). Climate change alters the future of natural floristic regions of deep evolutionary origins. Nat Commun, 15, 9474.

58. Montgomery, S.L. (1982). Biogeography of the Moth Genus Eupithecia in Oceania and the Evolution of Ambush Predation in Hawaiian Caterpillars (Lepidoptera: Geometridae). Entomologia Generalis, 8, 027–034.

59. Mougi, A. & Kondoh, M. (2012). Diversity of Interaction Types and Ecological Community Stability. Science, 337, 349–351.

60. Neu, A., Beaulieu, M. & Fischer, K. (2021). Limits on optimal decision making: host plant selection is not altered by high temperatures in a butterfly. Animal Behaviour, 174, 87– 95.

61. New, T.R. (2008). Insect conservation on islands: setting the scene and defining the needs. J Insect Conserv, 12, 197–204.

62. Nishida, R. (2002). Sequestration of Defensive Substances from Plants by Lepidoptera. Annual Review of Entomology, 47, 57–92.

63. Nylin, S. (1988). Host Plant Specialization and Seasonality in a Polyphagous Butterfly, Polygonia C-Album (Nymphalidae). Oikos, 53, 381–386.

64. Papaj, D.R., Mallory, H.S. & Heinz, C.A. (2007). Extreme weather change and the dynamics of oviposition behavior in the pipevine swallowtail, Battus philenor. Oecologia, 152, 365– 375.

65. Petrocelli, I., Alzate, A., Zizka, A. & Onstein, R.E. (2024). Dispersal-related plant traits are associated with range size in the Atlantic Forest. Diversity and Distributions, 30, e13856.

66. Revell, L.J. (2012). phytools: an R package for phylogenetic comparative biology (and other things). Methods in Ecology and Evolution, 3, 217–223.

67. Robinson, G.S., Ackery, P.R., Kitching, I., Beccaloni, G.W. & Hernández, L.M. (2023). HOSTS - a Database of the World’s Lepidopteran Hostplants.

68. Rubinoff, D. & Haines, W.P. (2005). Web-Spinning Caterpillar Stalks Snails. Science, 309, 575– 575.

69. Running, S., Mu, Q. & Zhao, M. (2021). MODIS/Terra Gross Primary Productivity 8-Day L4 Global 500m SIN Grid V061.

70. Schemske, D.W., Mittelbach, G.G., Cornell, H.V., Sobel, J.M. & Roy, K. (2009). Is There a Latitudinal Gradient in the Importance of Biotic Interactions? Annual Review of Ecology, Evolution, and Systematics, 40, 245–269.

71. Seastedt, T.R. & Crossley, D.A., Jr. (1984). The Influence of Arthropods on Ecosystems. BioScience, 34, 157–161.

72. Sessa, E.B., Chambers, S.M., Li, D., Trotta, L., Endara, L., Burleigh, J.G., et al. (2018). Community assembly of the ferns of Florida. American Journal of Botany, 105, 549–564.

73. Shapiro, A.M. & Cardé, R.T. (1970). HABITAT SELECTION AND COMPETITION AMONG SIBLING SPECIES OF SATYRID BUTTERFLIES. Evolution, 24, 48–54.

74. Shirey, V., Larsen, E., Doherty, A., Kim, C.A., Al-Sulaiman, F.T., Hinolan, J.D., et al. (2022) LepTraits 1.0 A globally comprehensive dataset of butterfly traits. Sci Data, 9, 382

75. Singer, M.S., Lichter-Marck, I.H., Farkas, T.E., Aaron, E., Whitney, K.D. & Mooney, K.A. (2014). Herbivore diet breadth mediates the cascading effects of carnivores in food webs. Proceedings of the National Academy of Sciences, 111, 9521–9526.

76. Skeels, A., Bach, W., Hagen, O., Jetz, W. & Pellissier, L. (2023). Temperature-Dependent Evolutionary Speed Shapes the Evolution of Biodiversity Patterns Across Tetrapod Radiations. Systematic Biology, 72, 341–356.

77. Slatyer, R.A., Hirst, M. & Sexton, J.P. (2013). Niche breadth predicts geographical range size: a general ecological pattern. Ecology Letters, 16, 1104–1114.

78. Storch, D., Bohdalková, E. & Okie, J. (2018). The more-individuals hypothesis revisited: the role of community abundance in species richness regulation and the productivity–diversity relationship. Ecology Letters, 21, 920–937.

79. Sun, L., He, Y., Cao, M., Wang, X., Zhou, X., Yang, J., et al. (2024). Tree phytochemical diversity and herbivory are higher in the tropics. Nat Ecol Evol, 8, 1426–1436.

80. Sunday, J.M., Bates, A.E. & Dulvy, N.K. (2010). Global analysis of thermal tolerance and latitude in ectotherms. Proceedings of the Royal Society B: Biological Sciences, 278, 1823–1830.

81. Townsend, J. & DiMiceli, C. (2015). MOD44B MODIS/Terra Vegetation Continuous Fields Yearly L3 Global 500m SIN Grid.

82. Wang, X., Luo, M., Song, F., Wu, S., Chen, Y.D. & Zhang, W. (2024). Precipitation Seasonality Amplifies as Earth Warms. Geophysical Research Letters, 51, e2024GL109132.

83. Yang, L.H. & Gratton, C. (2014). Insects as drivers of ecosystem processes. *Current Opinion in Insect Science*, Ecology, 2, 26–32.

84. Yotoko, K.S.C., Prado, P.I., Russo, C.A.M. & Solferini, V.N. (2005). Testing the trend towards specialization in herbivore–host plant associations using a molecular phylogeny of *Tomoplagia* (Diptera: Tephritidae). Molecular Phylogenetics and Evolution, 35, 701–711.

85. Zografou, K., Swartz, M.T., Tilden, V.P., McKinney, E.N., Eckenrode, J.A. & Sewall, B.J. (2020). Stable generalist species anchor a dynamic pollination network. Ecosphere, 11, e03225.

